# sCIN: A Contrastive Learning Framework for single-cell Multi-omics Data Integration

**DOI:** 10.1101/2025.02.03.636095

**Authors:** Amir Ebrahimi, Alireza Fotuhi Siahpirani, Hesam Montazeri

## Abstract

The rapid advancement of single-cell omics technologies such as scRNA-seq and scATAC-seq has transformed our understanding of cellular heterogeneity and regulatory mechanisms. However, integrating these data types remains challenging due to distributional discrepancies and distinct feature spaces. To address this, we present a novel single-cell Contrastive INtegration framework (sCIN), that integrates different omics modalities into a shared low-dimensional latent space. sCIN uses modality-specific encoders and contrastive learning to generate latent representations for each modality, aligning cells across modalities and removing technology-specific biases. The framework was designed to rigorously prevent data leakage between training and testing, and was extensively evaluated on three real-world paired and unpaired datasets including SHARE-seq, 10X PBMC (10k version), and CITE-seq. Paired datasets refer to multi-omics data generated using technologies capable of capturing different omics features from the same individual cells whereas unpaired datasets are collected from different but similar cell populations within the same tissue, with each modality measured separately. Results on paired and unpaired datasets show that sCIN outperforms state-of-the-art models, including scGLUE, scBridge, sciCAN, Con-AAE, Harmony, and MOFA+, across multiple metrics: Average Silhouette Width (ASW) for clustering quality, Recall@k, Cell type@k, Cell type accuracy, and Median Rank for integration quality. Moreover, sCIN was evaluated on simulated unpaired datasets derived from paired data, demonstrating its ability to leverage available biological information for effective multimodal integration. In summary, sCIN reliably integrates omics modalities while preserving biological meaning in both paired and unpaired settings.

## Introduction

The advent of single-cell omics technologies has transformed the study of biological systems, offering insights into phenotypes at single-cell resolution across different omics layers and cell types. For instance, single-cell RNA sequencing (scRNA-seq) quantifies gene expression by measuring messenger ribonucleic acid (mRNA) abundance in large numbers of individual cells [1]. Moreover, single-cell assay for transposase-accessible chromatin with high throughput sequencing (scATAC-seq) identifies open chromatin regions, which are essential for gene regulation due to the increased activity of transcription factors (TFs) and other regulatory elements in these regions [2]. Beyond scATAC-seq technologies, mapping chromatin accessibility [3–5] and DNA methylation patterns [6,7] further illuminate the molecular mechanisms underlying gene regulation.

Gene expression involves several stages, including transcription, post-transcriptional regulation, translation, and post-translational modifications. To better understand these processes, there is increasing demand for sequencing technologies that can measure multiple molecular features within a single cell. These multi-omics approaches offer a more holistic view of cellular function and help mitigate batch effects from separate experiments. For instance, SHARE-seq [8] and SNARE-seq [9] simultaneously profile the transcriptome and chromatin accessibility of a cell population by integrating DNA fragmentation and mRNA reverse transcription. Cellular Indexing of Transcriptomes and Epitopes by Sequencing (CITE-seq) [10] combines antibody-based tagging with reverse transcription to capture both transcriptome and cell surface protein profiles within the same cells. Analyzing each data modality independently can result in fragmented insights and an incomplete understanding of the biological system. Therefore, computational approaches have been proposed to integrate multiple omics data types for more comprehensive analysis and inference. However, this integration task is challenging due to disparities in the distributions and feature spaces of the different modalities.

Most computational approaches aim to learn a low-dimensional joint representation of multiple modalities using dimensionality reduction techniques such as Principal Component Analysis (PCA) [11], Canonical Correlation Analysis (CCA) [12–14], or deep neural networks [15–17]. For instance, Seurat [18] applies PCA to transform data linearly and uses Mutual Nearest Neighbors and CCA to align embeddings in a shared latent space. However, this approach is limited in capturing non-linear relationships between different omics features. Another approach, MOFA [19], is a probabilistic Bayesian framework that employs matrix factorization to decompose input data matrices from different modalities into two components: i) a modality-specific weight matrix that captures factor loadings for each feature, and ii) a shared factor matrix representing the factors loadings for each sample, which is common for all modalities. Harmony [20], on the other hand, uses fuzzy K-Nearest Neighbor (KNN) clustering to group cells based on cell types, while maximizing batch diversity within each cluster, offering a robust method for batch effect correction and modality alignment.

Many multimodal deep learning frameworks have also been developed to map different modalities’ high-dimensional, non-linear feature spaces into a unified, smaller subspace. For instance, scGLUE employs a graph Variational Autoencoder (VAE) to learn biological feature representations [17]. It enhances the cell embeddings learned by modality-specific VAEs using representations learned by the graph VAE. Additionally, adversarial training is applied to integrate technology effects, so an adversarial classifier cannot distinguish cell embeddings based on omics modalities. GCN-SC leverages mutual nearest neighbors to connect cells across datasets and construct a mixed graph. It also adjusts query datasets using a Graph Convolutional Network (GCN), followed by non-negative matrix factorization for dimension reduction and visualization [21]. MultiVI is a VAE-based framework that integrates scRNA-seq and scATAC-seq data by learning modality-specific latent representations using deep neural network encoders. It corrects batch effects and aligns these representations within a shared latent space [22]. Con-AAE uses adversarial Autoencoders to learn latent representations of paired single-cell transcriptome and chromatin accessibility data. By applying contrastive learning and a cycle consistency loss, it effectively integrates different data types and accounts for technology effects [23].

Here, we introduce sCIN, a contrastive learning-based neural network framework designed to integrate single-cell multi-omics datasets into a shared low-dimensional latent space. sCIN preserves cell type heterogeneity, eliminates technology effects, and applies to both paired and unpaired datasets. In particular, we adopt the CLIP model [24], which was originally designed to learn joint representations of images and text, to single-cell data. sCIN consists of two modality-specific encoders that learn reduced-dimensional representations for each omics type. The model then minimizes the distance between cells of the same type in latent space, while maximizing the distance between cells of different types through a contrastive loss function. We evaluated sCIN on four real-world paired and unpaired datasets: Ma-2020 (SHARE-seq) [9], 10x Genomics PBMC (10k version) [25], Luecken-2022 (CITE-seq) [26], and Muto-2021 (scRNA-seq and scATAC-seq of human kidney) [27]. We also simulated realistic, leakage-free unpaired datasets derived from these paired datasets to assess sCIN’s performance. The results from both settings show that sCIN outperforms six state-of-the-art and one baseline methods using various integration strategies across different evaluation metrics.

## Results

### Overview of sCIN

sCIN is a contrastive learning-based framework, inspired by CLIP [24], designed for integrating paired and unpaired single-cell multi-omics datasets. It uses two neural network encoders to map each data modality into a shared lower-dimensional space. For paired data, sCIN aligns embeddings by treating measurements from the same cell as positive pairs. For unpaired data, cells of the same type across modalities are considered positive pairs. Additionally, the model separates embeddings of cells with different cell types (negative pairs), ensuring that clustering reflects biological variability rather than technical biases from single-cell omics technologies. Integration quality is evaluated using metrics such as Recall@k, ASW, cell type accuracy, and median rank, and cell type@k (Methods). Recall@k and median rank assess the alignment of the same cells across modalities, while cell type accuracy, Average Silhouette Width (ASW), and cell type@k evaluate cell type consistency among nearest neighbors and clustering, respectively.

### Model evaluations on paired datasets

We benchmarked sCIN against six state-of-the-art methods: scGLUE[18], scBridge [28], sciCAN [29], MOFA+ [30], Con-AAE [23], and Harmony [20], as well as a baseline Autoencoder neural network trained solely with a reconstruction loss on three different paired datasets. All evaluations used ten train-test splits, resulting in ten hold-out test datasets. For metrics computed on separate embeddings, such as Recall@k, median rank, cell type accuracy, and cell type@k, we evaluated each metric using embeddings from the hold-out datasets across ten replications, resulting in ten values per metric for each model. We summarized these results by either plotting the mean with error bars (for Recall@k and cell type@k) or using box plots to illustrate variability across replications (for median rank and cell type accuracy).

### Evaluation of sCIN on the SHARE-seq Dataset

We evaluated sCIN on the SHARE-seq (Simultaneous High-throughput ATAC and RNA Expression with Sequencing) dataset, which profiles 32,231 mouse skin cells spanning 22 cell types. SHARE-seq enables the simultaneous measurement of gene expression and chromatin accessibility in single cells, offering a scalable and cost-effective approach [8]. The datasets contain 21,478 gene expression features and 340,341 chromatin accessibility features. The datasets were downloaded as h5ad files provided by Cao et al. [17]

We evaluated the models using the Recall@k metric, which measures the proportion of embeddings in the first modality whose corresponding counterparts from the second modality are among their k-nearest neighbors (Methods). On the SHARE-seq dataset, sCIN significantly outperformed other methods and achieved the highest Recall@k values across various k settings, with performance improving as *k* increased. Con-AAE also showed strong performance compared to other benchmarks, highlighting the potential of neural network-based frameworks for integration tasks while preserving biological information (Figure 2a).

**Figure 1.**
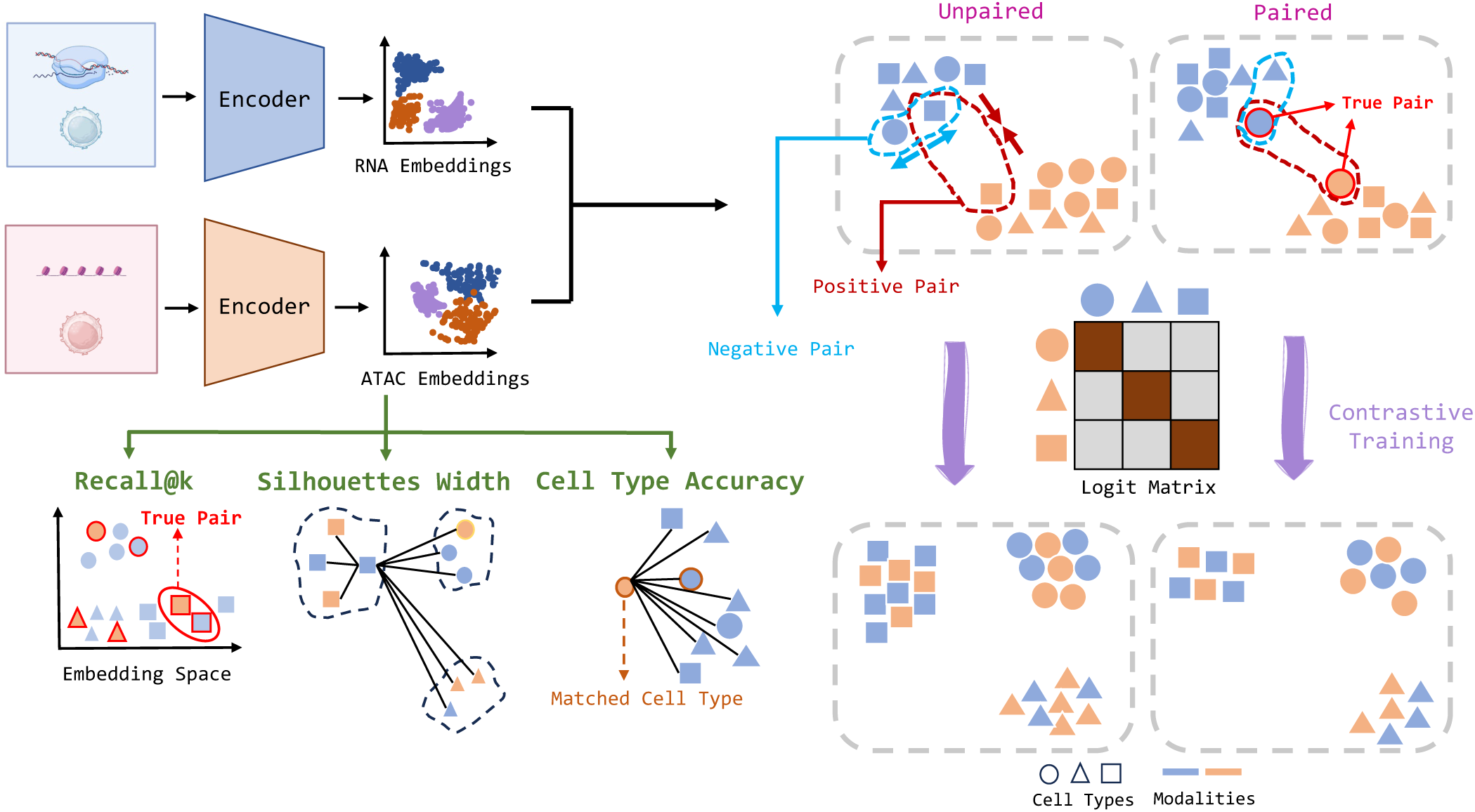
sCIN workflow. The framework uses modality-specific encoders to learn latent representations for each modality, using a contrastive training approach to align these representations. In the unpaired setting, cells of the same type are treated as positive pairs and are positioned closer together in the latent space. In contrast, cells of different types are treated as negative pairs and are separated further apart. In the paired setting, true cell pairs across modalities are defined as positive pairs. Performance is assessed using hold-out integrated embeddings, evaluated based on integration quality metrics (Recall@k) and the preservation of biological variability (ASW and cell type accuracy). Some parts of this figure were created using BioRender.com.

**Figure 2.**
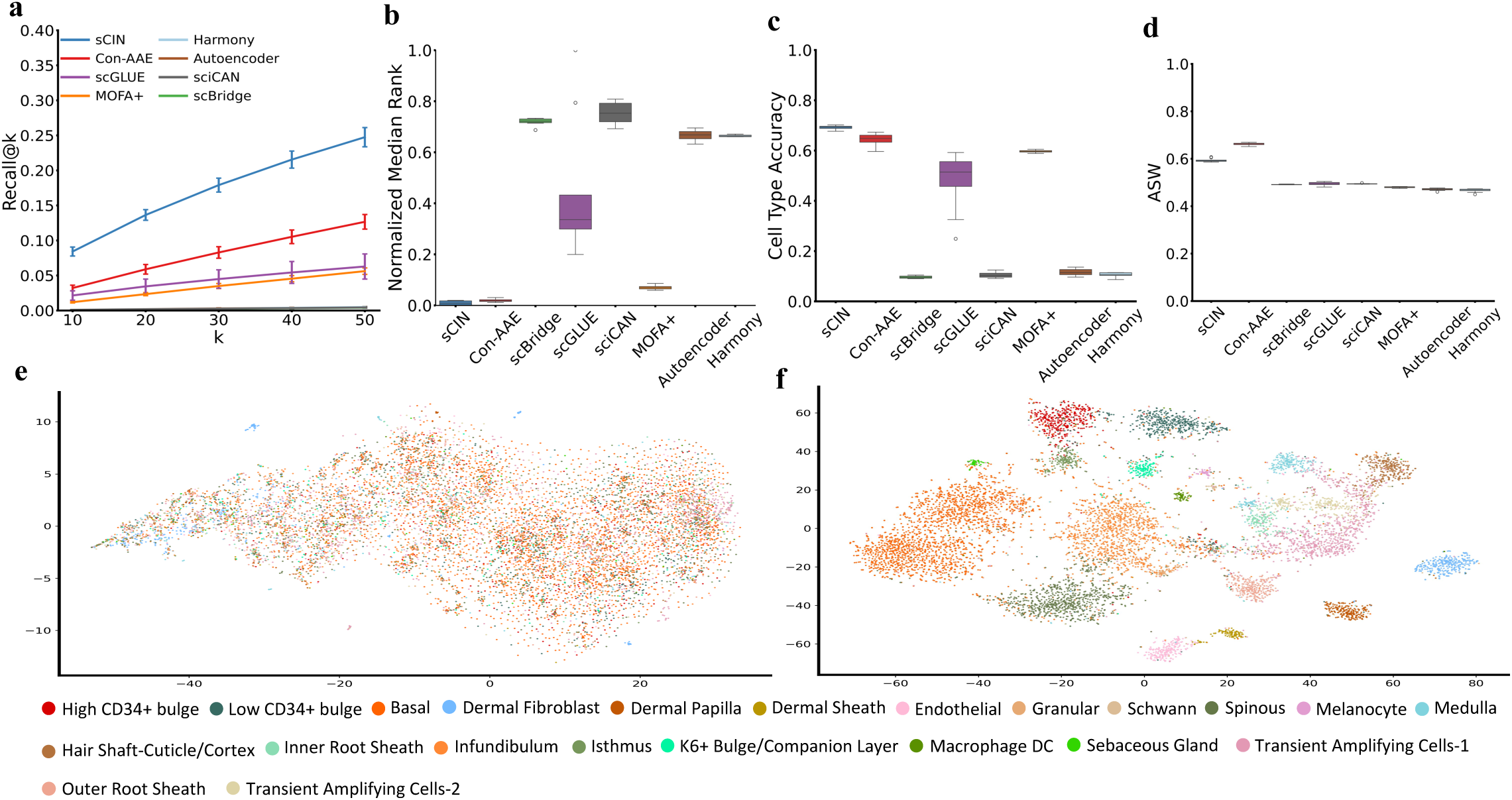
**a)** Comparison of Recall@k metrics between different models for k = 10, 20, 30, 40, and 50 for the SHARE-seq dataset. **b)** Normalized Median Rank values for different models. **c)** Cell type accuracy comparison between models. **d)** Comparison of ASW score across models. **e)** t-SNE visualization of the original data. **f)** t-SNE visualization of the sCIN’s integrated embeddings.

We used the Median Rank metric to assess data integration quality. For each cell, the distance between its embedding in the first modality and all embeddings in the second modality was calculated. The rank of the true pair’s distance among these values was obtained. The overall Median Rank was determined as the median of these ranks across all cells, with normalized values reported (Methods). On the SHARE-seq dataset, sCIN exhibited the closest pairwise distances in its embedding space compared to other models (Figure 2b). Consistent with the Recall@k results, the second and third best-performing models were Con-AAE and MOFA+, respectively.

We evaluated whether sCIN preserves cell type information post-integration, an aspect of biological interpretability in multi-omics integration. Cell type accuracy, assessed by the alignment of cell types among nearest neighbors in the embedding space (Figure 2c, d), showed that sCIN achieved the highest accuracy, nearly 0.7, outperforming all other models. Clustering quality, measured using ASW (Methods), ranked sCIN as the second-best model, with a normalized ASW exceeding 0.6, trailing Con-AAE by about 0.05.

To demonstrate that sCIN’s embedding space captures more relevant information than the original data, we visualized t-SNE representations of both the original data and the joint embeddings. To handle the high-dimensional nature of the multi-omics feature spaces, we first reduced the original features using PCA before applying t-SNE (Figure 2e). The t-SNE of the joint embeddings (Figure 2f), colored by cell types, shows better preservation of cell type information. On the SHARE-seq dataset, sCIN effectively preserved cell type information post-integration, while the original data’s t-SNE visualization captures less cell type information, likely due to noise and inherent sparsity in single-cell multi-omics data.

### Evaluation of sCIN on the PBMC Dataset

We conducted similar evaluations on the PBMC dataset (10k version), which includes paired profiling of gene expression and chromatin accessibility for peripheral blood mononuclear cells. This dataset was generated using the 10x Genomics sequencing platform and is widely used for benchmarking multi-omics integration methods [24]. The dataset comprised 9631 cells and 19 cell types, with 29095 genes and 107194 chromatin regions. The datasets were obtained as h5ad files provided by Cao *et al*.[17].

sCIN consistently achieved the highest Recall@k values across all k settings (Figure 3a). It demonstrated a near-zero normalized Median Rank, highlighting the closeness of embeddings of the two modalities in the embedding space (Figure 3b). The patterns for cell type accuracy and ASW were consistent with those observed in the SHARE-seq dataset (Figure 3c, d). Moreover, sCIN effectively retained most cell type information for 19 distinct cell types in the integrated embedding space (Figure 3e, f).

**Figure 3.**
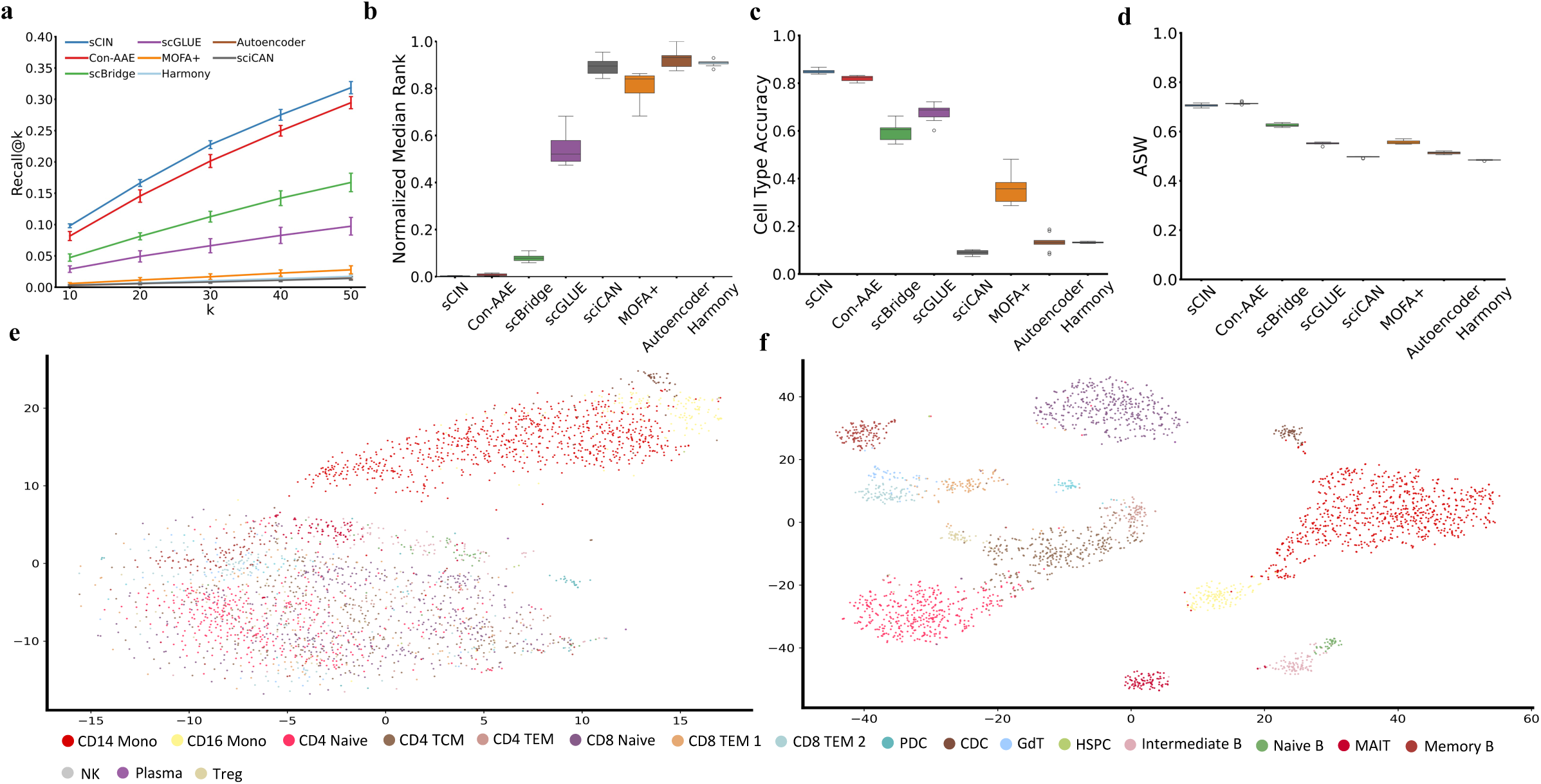
**a)**The comparison of Recall@k metrics between different models for k = 10, 20, 30, 40, and 50 for the 10X PBMC Multiome dataset (gene expression and chromatin accessibility). **b)** Normalized Median Rank values for different models. **c)** Cell type accuracy comparison between models. **d)** Comparison of ASW score across models. **e)** t-SNE embeddings of the original dataset. **f)** t-SNE visualization of the sCIN’s integrated embeddings.

### Evaluation of sCIN on the CITE-seq Dataset

To show sCIN’s ability to handle diverse omics modalities, we evaluated its performance on the CITE-seq dataset [26], which includes gene expression and cell surface protein features. Single-cell CITE-seq contains 90,261 matched scRNA-seq and ADT profiling [10] of bone marrow mononuclear cells of 12 healthy donors across different sites. The data was generated using the 10x 3’ Single-Cell Gene Expression kit with Feature Barcoding and the BioLegend TotalSeq B Universal Human Panel v1.0. It includes measurements of 13,953 genes and 134 cell surface proteins. The datasets were downloaded from the GEO accession number GSE194122 as h5ad files.

On this dataset, sCIN outperformed all other models in integrating quality and preserving biological information. Specifically, sCIN consistently increased Recall@k values as k increased, while other frameworks plateaued and showed no significant improvements (Figure 4a). A similar trend was observed in the Median Rank metric (Figure 4b), where sCIN achieved the closest alignment of cell pairs across modalities. Additionally, sCIN achieved the highest performance in cell type accuracy and ASW metrics, outperforming other methods (Figures 4c, d).

**Figure 4.**
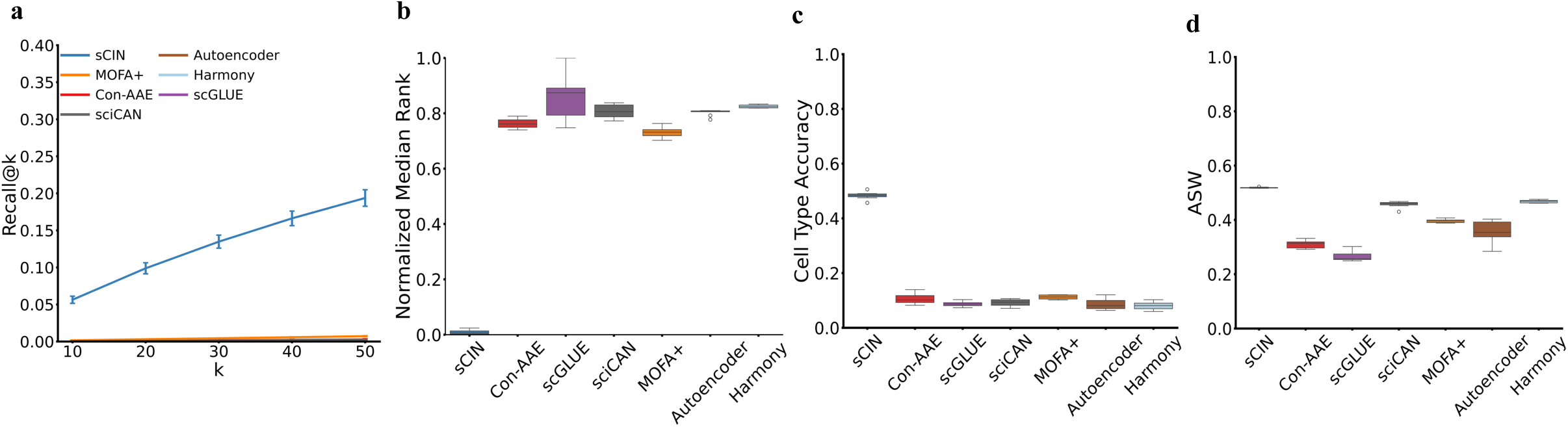
**a)** The comparison of Recall@k metric between different models for k = 10, 20, 30, 40, and 50 for the CITE-seq dataset (gene expression and cell surface proteins). **b)** Normalized Median Rank metric for different models. **c)** Cell type accuracies between models that show the percentage of closest embeddings from different modalities having the same cell types. **d)** Comparison of ASW score across models shows the clusters’ quality.

### Improved Cell Type Clustering with sCIN’s Joint Embeddings

To compare sCIN’s latent space with the original and PCA-transformed spaces, we computed ASW for the original data modalities, joint PCA embeddings, and sCIN’s joint embeddings. PCA dimensionality matched sCIN’s (256 for all datasets). Results show sCIN’s embeddings achieve superior cell type clustering over PCA and original data. This experiment was performed on test datasets and replicated 10 times (Supplementary Figure 1).

### Model evaluations on simulated unpaired datasets

We assessed the performance of sCIN on unpaired datasets, where omics data from different modalities do not originate from the same set of cells. First, we simulate unpaired datasets from the paired datasets: SHARE-seq, PBMC, and CITE-seq. Specifically, a subset of cells was selected, and only one modality (ATAC for SHARE-seq and PBMC, and ADT for CITE-seq) from this subset was retained. For the remaining cells, only the data for the second modality was used, ensuring no overlap between the cells of the two modalities. To examine the impact of unbalanced cell proportions across modalities, we generated datasets with varying levels of proportions (1%, 5%, 10%, 20%, and 50%) and compared the resulting embeddings on different metrics: Recall@k, ASW, and cell type accuracy. We also included the paired datasets as the optimal case and a random baseline, where cell types were randomly permuted and assigned to cells (Figure 5). All evaluations were performed on the hold-out test datasets and replicated 10 times.

**Figure 5.**
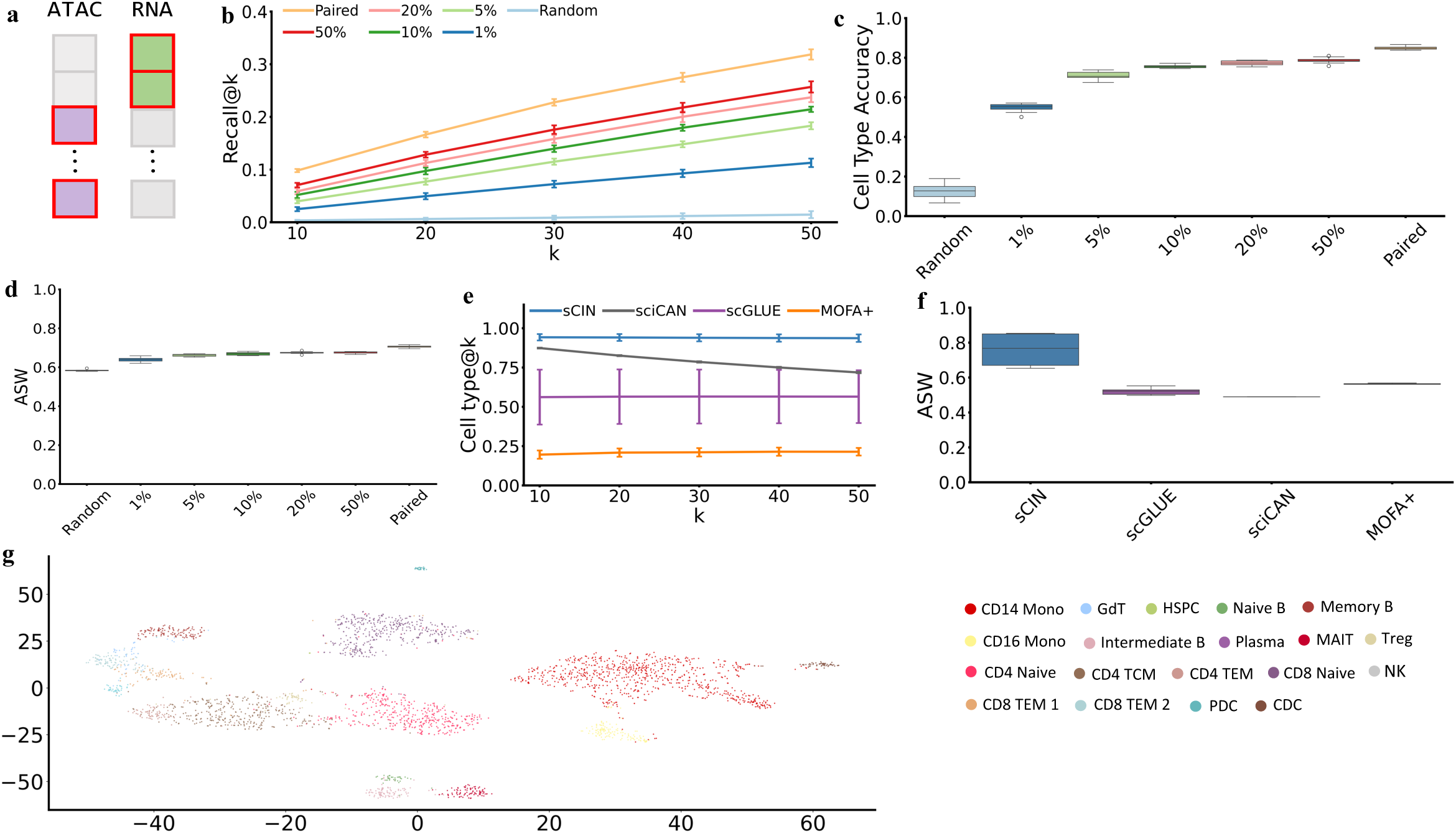
Evaluation of sCIN on unpaired datasets. **a)** Schematic showing the simulation of unpaired data from paired data. **b)** Recall@k performance on the simulated unpaired datasets. **c)** Cell type accuracy performance on the simulated unpaired datasets. **d)** ASW based on joint embeddings from the simulated unpaired datasets. **e)** Comparison of sCIN and the state-of-the-art models based on the cell type@k metric on the Muto *et al.* dataset. **h)** Comparison of sCIN and the state-of-the-art models based on the ASW metric on the Muto *et al.* dataset. **g)** t-SNE representations of the embeddings from the hold-out simulated unpaired PBMC dataset colored by cell types.

On the PBMC dataset, Recall@k values significantly improved as the proportion of cells for each cell type increased. Although the results remained lower than the paired case, they were significantly higher than the random baseline (Figure 5b). In the cell type accuracy analysis, as fewer cells were provided for the second modality, the cell type accuracy decreased. The integrated embeddings from the paired dataset achieved the highest accuracy (0.848 on average). Across proportions from 1% to 50%, accuracy showed an increasing trend, consistently outperforming the random baseline (Figure 5c).

Additionally, we assessed the ASW score of the embeddings, demonstrating the preservation of cell type information (Figure 5d). To further assess robustness, we calculated Recall@k and cell type accuracy from gene expression embeddings to chromatin accessibility embeddings (Supplementary Figures 4ab); previous results showed similar results when comparing chromatin accessibility to gene expression. We also visualized t-SNE representations of the hold-out PBMC dataset in the embedding space of sCIN trained on 50% unpaired data (Figure 5g), which clearly shows the clustering of integrated embeddings based on cell types. Similar trends were observed in experiments using the the SHARE-seq dataset (supplementary Figure 2a and supplementary Figure 4bc) and on the CITE-seq dataset (supplementary Figure 3 and supplementary Figure 4ef).

### Model evaluations on Muto-2021 dataset

We further evaluated sCIN using real-world, unpaired single nucleus RNA sequencing (snRNA-seq) and single nucleus ATAC sequencing (snATAC-seq) data from Muto *et al*. [27]. The datasets were downloaded as h5ad files provided by Cao *et al*. [17]. The gene expression dataset consists of 19985 cells and 27,146 genes while the chromatin accessibility dataset has 24,205 cells and 99,019 open chromatin regions (i.e., peaks). These datasets obtained from non-tumor kidney cortex samples from patients undergoing partial or radical nephrectomy for renal mass. snRNA-seq and snATAC-seq libraries were obtained using 10X Genomics Chromium Single Cell 5′ v2 and 10X Genomics Chromium Single Cell ATAC v1, respectively. Because the datasets are unpaired—with no direct correspondence between cells across modalities—we focused on evaluation metrics that capture cell type information in the learned embeddings. Specifically, sCIN consistently showed high performance across all *k* values (10, 20, 30, 40, 50) (Figure 5e), followed by sciCAN and scGLUE. Similar results were obtained in the ASW evaluation (Figure 5f).

## Methods

### sCIN framework

sCIN is a contrastive learning-based framework designed to learn a shared low-dimensional latent embedding across modalities of single-cell data. sCIN is developed based on the CLIP architecture developed by OpenAI, which aligns image and text embeddings in a shared latent space. sCIN extends this approach to integrates single-cell modalities such as scRNA, scATAC, and single-cell ADT profiles. It supports both paired and unpaired datasets: in the paired case, the input consists of simultaneous measurements of the same single cells across two modalities, whereas in the unpaired case, the two modalities originate from different single cells.

We used two neural network encoders to map the distinct feature spaces of each modality into a shared latent space. The main objective is to align the latent embeddings of each cell across modalities, irrespective of the technology used. sCIN can also handle unpaired datasets, where each cell is measured in only one modality. In this case, sCIN aims to bring embeddings of cells from the same cell type closer together while separating those from different cell types.

Let (*x*_*i*_ *y*_*i*_) for *i* = 1…*N* denote the raw feature values for the first and second modalities, where *N* is the number of cells, and *x*[*i*] ∈ *R*^*p*^ and *y*[*i*] ∈ *R*^*q*^. Here, *p* and *q* represent the number of features for the first and second modalities, respectively. We used two encoders, *f* and *g*, for the first and second modalities to generate latent embeddings for both modalities, each with the same dimension size *d*. The embeddings are normalized such that ||*f*()|| = 1 and ||*g*()|| = 1.

In a contrastive learning framework, positive and negative pairs are defined to guide the algorithm in learning embeddings where positive pairs are closer and negative pairs are further apart. In the paired case, the pair (*x*[*i*] *y*[*i*]) is considered a positive pair, while (*x*[*i*] *y*[*j*]), where cells *i* and *j* are from different types, is considered a negative pair. Note that (*x*[*i*] *y*[*i*]) where *i* and *j* are distinct cells *i* and *j* of the same type, are neither a positive nor negative pair. In the unpaired case, (*x*[*i*] *y*[*j*]), where cells *i* and *j* share the same type, are treated as positive, while all others are treated as negative pairs.

sCIN constructs *M* × *M* matrices for mini-batches, consisting of similarity between embedding of the first and second modalities, where *M* represents the number of cell types (Figure 1). The diagonal entries represent positive pairs, while off-diagonal entries represent negative pairs. sCIN learns a shared embedding space by jointly training the encoders to maximize the similarity of diagonal entries while minimizing the off-diagonal pairs. The loss function for a given anchor cell *i* is given as follows,

The total loss is summed up across all anchor cells in the mini-batch. In the unpaired case, the loss function is similar; however, the similarity matrix is constructed using a set of *M* cells from different types in the first modality and a corresponding set of *M* cells from the second modality, whereas, in the paired cases, the same set of cells is used for both modalities.

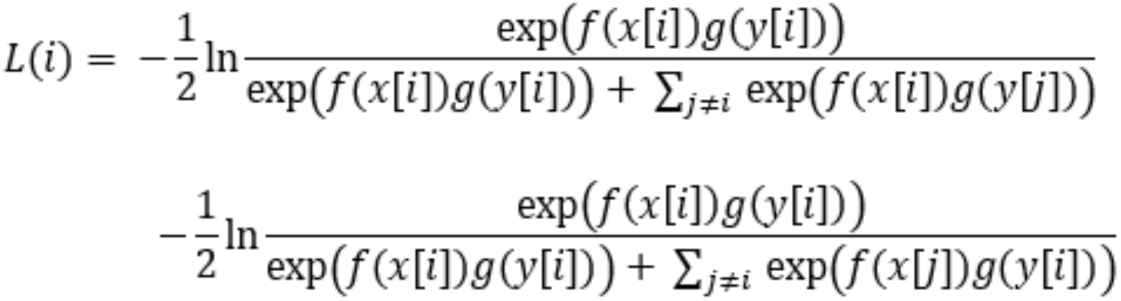

sCIN was trained in 150 epochs for all datasets, with a custom early-stopping strategy implemented to enhance training efficiency. The early stopping criteria used a patience of 10 and a minimum delta of 10^−4^. The encoders share the same architecture, consisting of three layers. The first linear layer projects the input data into a hidden dimension (256), followed by a batch normalization layer. A ReLU activation function is applied to the last linear layer to introduce non-linearity relationships between features. The final linear layer outputs latent embeddings of 128 dimensions.

### Benchmark models

To ensure rigorous evaluation and avoid any train-test data leakage, we implemented a unified modular training and evaluation framework for all models. This was particularly important as we identified instances of such leakage in some published models. Our framework strictly restricted the training function to access only the training data, excluding any information from the test set. The trained model was subsequently used to generate embeddings for the hold-out dataset, enabling fair and consistent evaluations across all models (see more details at https://github.com/AmirTEbi/sCIN). The embeddings generated by all models were evaluated using five metrics: ASW, Recall@K, and median rank, cell type accuracy, and cell type@k. Additionally, for each model, a joint embedding was constructed by concatenating embeddings from all modalities. sCIN was implemented and trained using the PyTorch framework in Python on the Nvidia GeForce RTX 4090 Graphical Processing Unit (GPU). Each benchmark experiment was replicated 10 times.

### Con-AAE

It uses two encoders to map each modality into low-dimensional manifolds, employing an adversarial loss to ensure that an adversarial classifier cannot distinguish between the latent embeddings based on their modalities [23]. To enhance stability, the model incorporates a cycle-consistency loss. Additionally, a contrastive learning loss is applied to the embeddings to increase similarity between cells of the same type. Finally, a classifier predicts cell type probabilities based on the joint embedding space. The model was trained using default parameters, and the original Con-AAE implementation is available at https://github.com/kakarotcq/Con-AAE/tree/main.

### MOFA+

Multi-Omics Factor Analysis v2 (MOFA+) [30] is an unsupervised, probabilistic matrix factorization framework designed for the integration of single-cell multi-omics data. It extends the original MOFA model [19] by introducing a more scalable variational-inference engine, flexible sparsity constraints, and explicit support for modeling multiple sample groups (“multi-group” functionality) while handling missing values across assays. Using the mofapy2 Python package (https://github.com/bioFAM/mofapy2), we trained MOFA+ on the entire dataset, as it lacks modality-specific embeddings and train/test splitting. To evaluate MOFA+’s paired test embeddings based on Recall@k, cell type accuracy, and median rank, we imputed one modality from another by augmenting the dataset with test samples initialized as zero matrices. For the unpaired test datasets, we extracted the learned weights, precisions, and intercepts for each view, and computed the closed-form posterior mean of the factors by centering and precision-weighting the test data matrix, and finally multiplied by the precomputed weight–covariance product. For efficiency, chromatin accessibility was reduced to 100 PCs before training. MOFA+ was trained with 1000 iterations, 30 factors, and “fast” convergence mode.

### Harmony

Harmony refines PCA embeddings through two steps: (1) diversity-maximizing clustering and (2) mixture model-based linear correction [20]. Although originally developed for batch correction, we adapted Harmony for multimodal integration. Datasets were reduced to 100 PCs and then projected into shared space using Canonical Correlation Analysis. Preprocessing followed harmonypy standards as in https://github.com/slowkow/harmonypy. Since harmonypy does not natively support train/test splits, we trained Harmony on the training set (10 epochs). For evaluation, we cloned the model with identical parameters to generate embeddings for the hold-out data in a single epoch.

### scGLUE

scGLUE consists of different modules [17]. It employs modality-specific VAEs to generate cell embeddings. Moreover, it encodes the relationships between biological features (e.g., genes and chromatin regions) in a bi-directional graph with self-loops. A graph VAE is then used to produce embeddings for each node (i.e., feature) in that graph. Since the latent dimensions of the cells and feature embeddings are aligned, scGLUE reconstructs the original data using the inner product of cells and feature embeddings. It also employs an adversarial classifier to predict the omics modality using cell embeddings. scGLUE was run using the ‘scglue’ python package based on the default settings and guidelines (https://scglue.readthedocs.io/en/latest/tutorials.html).

### scBridge

scBridge is a semi-supervised method for integration of paired scRNA-seq and scATAC-seq data [28]. scBridge uses annotated scRNA-seq data to find representative cell-type features (prototypes). Then, it accepts the scATAC-seq data as gene activities and identifies scATAC-seq cells showing high correlation between chromatin accessibility and gene expression of the prototypes as reliable candidates for integration. scBridge was run using the default settings as stated in https://github.com/XLearning-SCU/scBridge/tree/main.

### sciCAN

sciCAN (single-cell chromatin accessibility and gene expression data integration via Cycle-consistent Adversarial Network) is an unsupervised deep learning model that employs a shared encoder to learn integrated representations from both modalities, utilizing noise contrastive estimation to enhance discriminative features [29]. To align the modalities, sciCAN incorporates a cycle-consistent adversarial network, enabling the translation of data between modalities and ensuring consistency through cycle-consistency loss. Its architecture allows sciCAN to integrate data without requiring paired cells or prior annotations. sciCAN was run using the default settings according to https://github.com/rpmccordlab/sciCAN.

### Autoencoder

The baseline Autoencoder uses separate encoders for each data modality, mapping inputs to a 64-dimensional embedding via a 128-dimensional hidden layer with batch normalization and ReLU activation. These embeddings are concatenated and fed into a decoder to reconstruct the inputs. For the SHARE-seq and PBMC datasets, the hidden and latent dimensions were set to 256 and 128, respectively, with 2000, 500, and 100 PCs selected for SHARE-seq, PBMC, and CITE-seq. Training was conducted for 150 epochs with a learning rate of 0.01, and a batch size of 64 for all datasets.

### Evaluation metrics

#### ASW

Silhouette Width evaluates clustering quality by comparing a cell’s within-cluster distance to its distance from the nearest neighboring cluster. The ASW, calculated as the mean Silhouette Width across all cells, ranges from −1 to 1, with higher values indicating better cluster separation. To assess integration results, we used ASW based on cell type labels. Euclidean distance was employed as the distance metric. Specifically, embeddings generated by each method for different modalities were concatenated in feature space, and ASW was calculated on the integrated embeddings. We normalized ASW to a range between 0 and 1 as follows:

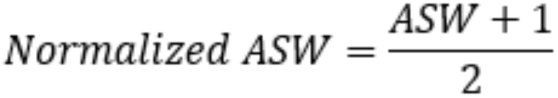

The higher value for this metric indicates the model’s ability to preserve cell-type heterogeneity in the embedding space effectively.

#### Recall@k

Recall@k metric assesses the closeness of different embeddings for the same cells in latent space, obtained using different modalities. To achieve this, the embedding of cell *i* from the first modality (e.g., RNA) is compared to all cell embeddings in the second modality (e.g., ATAC), and the *k*-nearest neighbors of cell *i* are then computed. We denote *N*^2^(*i*) as the cell indices of the k-nearest neighbors in the second modality. Irrespective of training on paired or simulated unpaired data,

since the hold-out data is paired, we can assess whether the matched embedding of the same cell is among the k-nearest neighbors. The fraction of such cells is called Recall@k and is computed as:

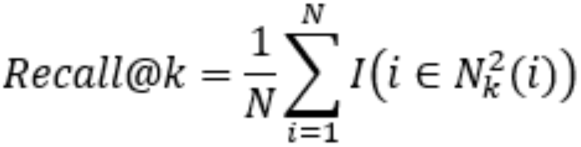

Where *I*() is an indicator function that equals 1 if cell *i* is among the k-nearest neighbors in the second modality, and 0 otherwise and *N* is the number of cells. Recall@k ranges from 0 to 1, with higher values indicating better matching of paired cell embeddings across modalities.

#### Cell type@k

This metric computes, for each cell in modality 1, the fraction of its *k* nearest neighbors (in the learned embedding space of modality 2) that share the same cell-type label, and then averages these proportions across all cells.

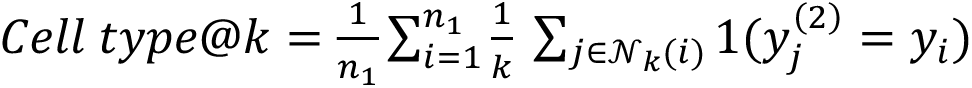

Where *n*_1_ denotes the number of cells in the modality 1, *N*_*k*_(*i*) represents the set of *k* nearest neighbors to cell *i* in the embedding space of modality 2, *y*_*i*_ is the cell type label of *i,* and *y_j_*^(2)^ is the cell type label of the *j*th neighbor in modality 2.

#### Cell type accuracy

The procedure for this metric is similar to Recall@k but instead checks whether the cell type of the query cell matches that of its one-nearest neighbor in the second modality.

#### Median Rank

For each cell embedding in the first modality, we calculated its Euclidean distances to all embeddings in the second modality and identified the rank of its matched embedding. The median rank across all cells serves as the metric, with lower values indicating better performance in accurately matching different data modalities for the same cells.

### Data Preparation

Preprocessed SHARE-seq, PBMC, and unpaired snRNA-seq and snATAC-seq data were downloaded from https://scglue.readthedocs.io/en/latest/data.html. Preprocessed CITE-seq data was downloaded from the Gene Expression Omnibus database with the accession number GSE194122.

## Conclusion

Contrastive Learning has emerged as a significant representation learning method in machine learning, specifically in image analysis and natural language processing [31,32]. It has also been applied in single-cell omics research for data analysis and integration [33,34]. The core idea of contrastive learning is to align semantically similar samples (positive pairs) closely in the embedding space while pushing dissimilar samples (negative pairs) further apart. Positive and negative pairs can be from different modalities such as natural language and images.

We introduced sCIN, a novel framework designed to effectively integrate paired and unpaired single-cell omics data modalities, including scRNA-seq, scATAC-seq, and scADT. sCIN leverages different omics from the same cells in paired datasets and from cells of the same cell types in unpaired datasets. Our method outperforms alternative approaches, such as *k*-nearest neighbor clustering, matrix factorization, and neural networks as well as baseline models (Autoencoder and PCA) in integrating multi-omics datasets while preserving essential biological characteristics, including cell type identity. Notably, sCIN achieved higher ASW scores than other methods, reflecting superior clustering quality based on cell types. Additionally, the comparison of sCIN’s ASW scores with those of the original datasets and the PCA baseline underscores its ability to extract meaningful biological insights from sparse single-cell data.

sCIN not only integrates paired multi-omics datasets but also effectively learns representations from unpaired omics data with different cell counts. Across various unpairing strategies, sCIN consistently captured biological information, with acceptable performance with limited sample sizes. Moreover, sCIN was designed to prevent data leakage between training and evaluation. The implementation is publicly available (Code Availability), accompanied by a tutorial for benchmarking other methods.

sCIN’s reliance on cell-type labels during training is a limitation due to the challenges of annotating single-cell omics data. Incorporating unsupervised or self-supervised learning approaches could address this issue. Additionally, the current framework is restricted to integrating two omics modalities. Future work could focus on extending the sCIN architecture to handle multiple data types and tackle specific biological tasks, such as gene regulatory network inference.

### Key Points

- sCIN is a novel contrastive learning-based framework for integrating paired and unpaired single-cell multi-omics datasets.
- sCIN consists of two modality-specific encoders, trained using a contrastive loss function tailored for single-cell integration tasks.
- sCIN outperformed alternative models based on matrix factorization, Fuzzy KNN clustering with linear batch correction, and neural network models in a rigorous assessment.

## Data availability

Preprocessed SHARE-seq, PBMC, and unpaired snRNA-seq and snATAC-seq datasets can be accessed from https://scglue.readthedocs.io/en/latest/data.html. The CITE-seq dataset is publicly available in the Gene Expression Omnibus (GEO) database with GSE194122 accession number.

## Code availability

Code for sCIN implementation and all analyses in this paper is publicly available at https://github.com/AmirTEbi/sCIN.

## Author contributions

A.E. and H.M. conceived the study and designed the experiments. A.E. and H.M. developed the method. A.E. collected and preprocessed the data, implemented the methods in Python, conducted all the experiments, analyzed the results, and wrote the manuscript. A.F.S. and H.M. supervised the project and verified the findings. A.E., H.M., and A.F.S. discussed the results and contributed to the final version of the manuscript. All authors agreed to the final version of the manuscript.

