## Supplementary Material for "sCIN: A Contrastive Learning Framework for single-cell Multi-omics Data Integration"

### Supplementary Figures

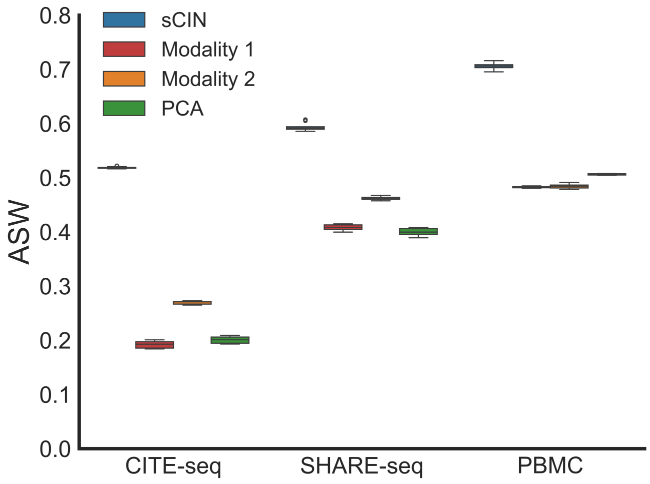


**Supplementary Figure 1.** Comparison of ASW metric between each modality, joint PCA embeddings of modalities, and joint sCIN embeddings for each paired dataset.

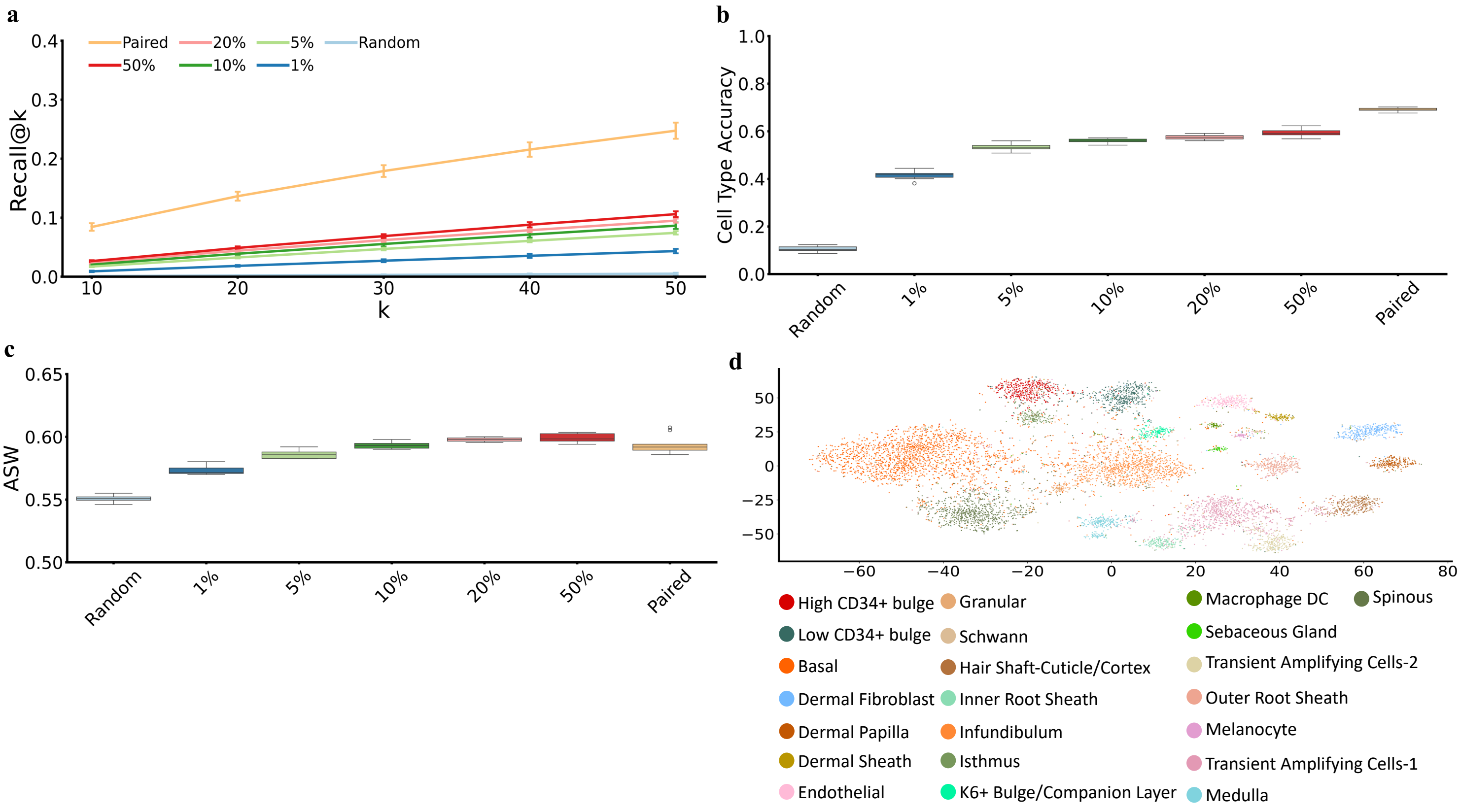


**Supplementary Figure 2.** Comparison of metrics across unpaired, random, and paired settings in the SHARE-seq dataset. **a)** Recall@k **b)** Cell type accuracy **c)** ASW based on the joint embeddings **d)** t-SNE representations of the embeddings from the hold-out dataset colored by cell types. delete

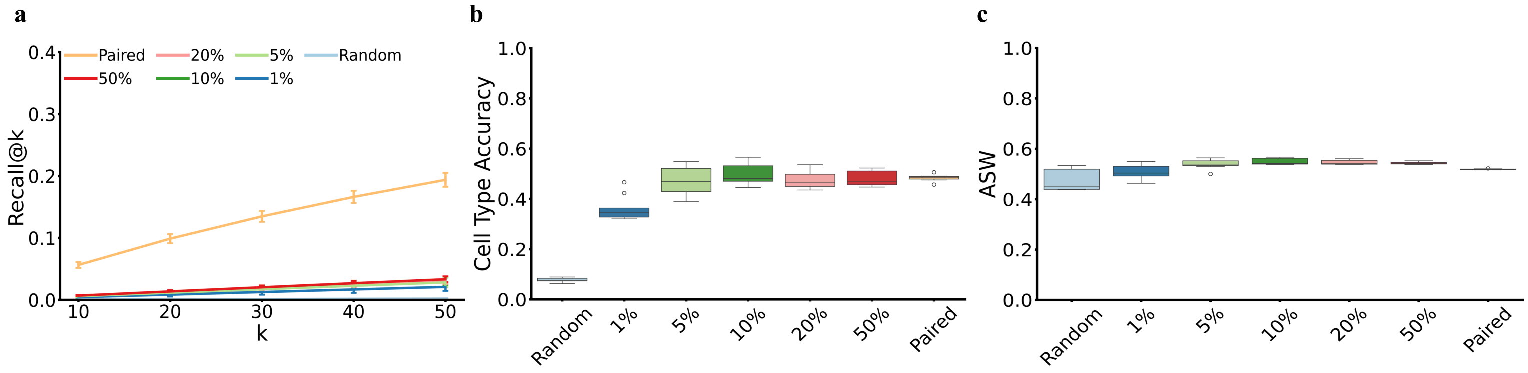


**Supplementary Figure 3.** Comparison of metrics across unpaired, random, and paired settings in the CITE-seq dataset. **a)** Recall@k **b)** Cell type accuracy **c)** ASW based on the joint embeddings

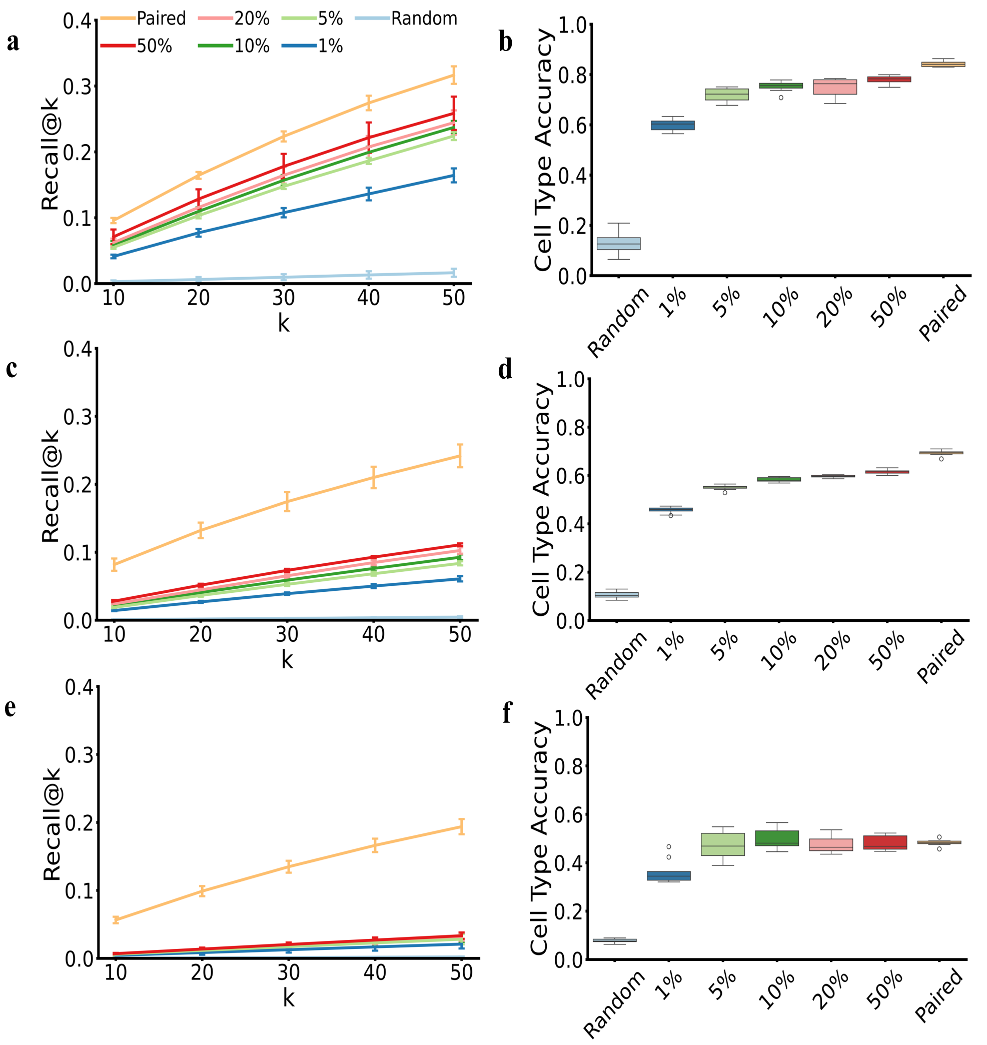


**Supplementary Figure 4.** Recall@k and cell type accuracy values from **a,b)** gene expression to chromatin accessibility for PBMC data, **c,d)** gene expression to chromatin accessibility for SHARE-seq data and **e,f)** gene expression to cell surface proteins for CITE-seq data.
